# Exploring the Energy Landscape of Hairpin Folding using the TIS-DNA model

**DOI:** 10.64898/2026.02.24.707746

**Authors:** Krishnakanth Baratam, Debayan Chakraborty

## Abstract

Coarse-grained (CG) modeling of DNA has been gathering steam in recent years due to the limitations imposed by current computer hardware, and deficiencies of atomistic force-fields. Specifically, CG simulations have emerged as a potent tool for exploring complex energy landscapes underlying biochemical processes, such as DNA transcription, as well as material design based on programmable self-assembly. In this chapter, we illustrate how the Three Interaction Site (TIS) model for DNA, a robust coarse-graining framework, can be used to study the folding landscape of DNA hairpins. We show that despite its simplicity, the TIS-DNA model quantitatively describes the hairpin folding thermodynamics and recapitulates many features of the kinetics, including the multiplicity of pathways. The free energy landscape exhibits ‘single-funnel’ character with a distinct bias towards the folded state. It is likely that folding initiates through non-specific collapse of the DNA chain, involving multiple excursions on the energy landscape, until the opposing strands are approximately aligned. Subsequently, the loop region becomes more ordered, and after the first native-contact nucleates, the rest of the process becomes essentially downhill.

## Introduction

DNA, ‘the blueprint of life’, performs a plethora of biological functions through highly complex motions spanning multiple length and timescales.^1^ While microscopic fluctuations at the base-pair level are crucial for protein recognition, several processes, including transcriptional regulation, are facilitated by mesoscale reorganization. ^2^ The tunable dynamics of DNA, which allow it to access various functional states in response to cellular or environmental cues, allude to a complex mechanism whose molecular details have not yet been fully resolved. To come up with with a reasonable answer, the intricate link between structure, dynamics and ultimately function must be understood at the molecular level. The theoretical underpinnings of these intimate connections are best described using the language of energy landscape theory.^3^

During the last few decades or so, innovations in experimental methodologies have provided crucial insights into DNA conformational landscapes. Ansari and coworkers in their insightful study^4^ showed that folding kinetics is strongly dependent on initial conditions, with temperature-jump and microfluidic experiments reporting rate constants that differ by an order of magnitude. Using Fluorescence Correlation Spectroscopy (FCS), Van Orden and coworkers illustrated that the folding process can be extremely slow (extending to *ms* time-scales and beyond), featuring long-lived intermediates.^5,6^ In another study, Breslauer and coworkers quantified the ruggedness of DNA landscapes using calorimetric and spectroscopic measurements,^7^ and alluded to its role both in the context of function and disease.

The emergence of single-molecule techniques has also been a game-changer. Several pioneering studies^8–11^ have revealed that the folding landscapes of even simple structural motifs, such as hairpins, could be quite complex, with multiple intermediates and kinetic traps, as well as a distribution of transition path times. These remarkable experimental findings have caused a paradigm shift in our understanding of DNA hairpin folding from a classical ‘two-state’ process^12,13^ to one exhibiting multi-state kinetics. For further details, we refer the reader to several authoritative reviews on this subject.^14,15^ It is worth emphasizing that despite DNA’s structural repertoire being less extensive than that of RNA, their folding mechanisms share several common themes, most notably kinetic partitioning and non-Arrhenius kinetics. ^16–18^

### Computational Models of DNA

Although several landscape parameters can be deduced from experiments alone, a reliable computational model is required to unravel the underlying microscopic features. While it is appealing to represent DNA, the surrounding as well as the counter-ions at the atomistic detail, such an approach is often not computationally feasible. The innumerable degrees of freedom coupled together makes the underlying energy landscape too complex (Figure 1D). Due to limitations in current hardware, traversing such high-dimensional landscapes to extract information over biologically relevant spatio-temporal scales remains practically intractable, particularly for complex DNA molecules. Strikingly, conventional molecular dynamics simulations based on atomistic force-fields have proved extremely challenging for probing the folding landscapes of even simple constructs, such as DNA triloop hairpins due to the time-scales involved.^19^ To mitigate this sampling bottleneck, an assortment of enhanced sampling techniques have been utilized. For instance, Zacharias and coworkers used replica exchange molecular dynamics (REMD) in conjunction with umbrella sampling to identify folding intermediates and quantify sequence-dependent energetics.^20,21^ However, in such enhanced sampling setups, temporal information of the folding process is lost due to the exchanges in temperature/Hamiltonian space and the addition of biasing potentials. In this context, the study by Portella and Orozco^22^ represents a landmark contribution, and is probably the first of its kind. Using unbiased molecular simulations, the authors unequivocally showed that hairpin folding is characterized by multiple routes, spans a range of time-scales, and involves long pauses in metastable states.

**Figure 1.**
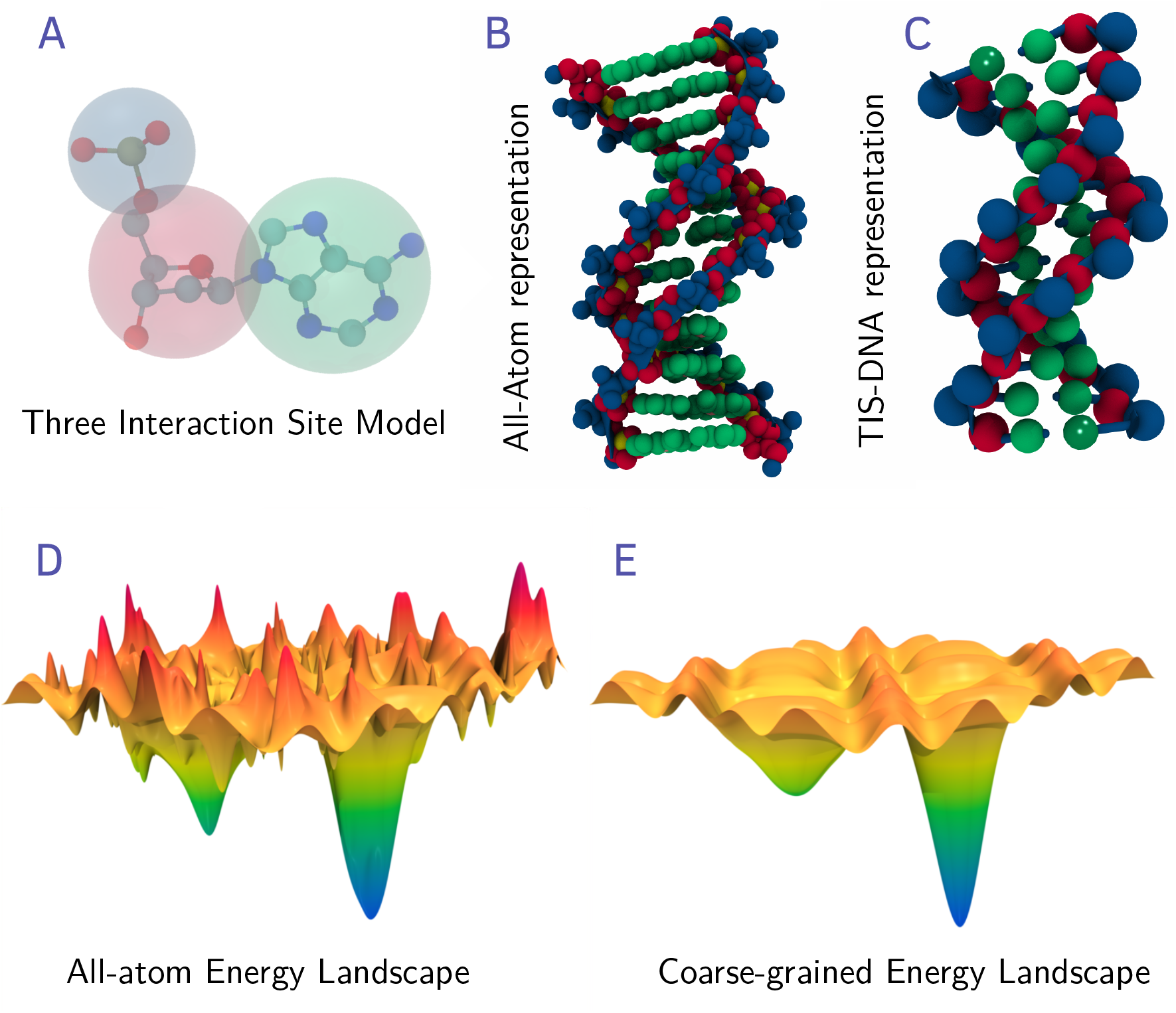
(A) The Three Interaction Site (TIS) scheme for grouping various atoms. The phosphate bead is colored blue, the sugar bead is colored red, and the nucleobase bead is colored green. (B) All-atom representation (left) of a double helix, and the corresponding TIS-DNA representation (right). (D-E) A schematic illustrating how coarse-graining drastically simplifies the underlying energy landscapes, by removing unnecessary features (‘ruggedness’), while still preserving the key features (relevant minima and saddle points) of the atomistic potential energy surface.

The concerted efforts of various groups to characterize DNA conformational dynamics using atomistic force-fields are indeed commendable, and have been highlighted in various recent reviews.^23,24^ But for many applications, it is often desirable to use a simplified or ‘coarse-grained’ (CG) description of DNA, with the level of coarse-graining dictated by the particulars of the problem. For example, understanding the complex organization of chromatin architecture typically requires representing about 1000 bp of DNA by a single interaction site,^25,26^ whereas deciphering the details of hairpin folding kinetics requires a more fine-grained description, with interaction sites centered on individual nucleotides. An immediate consequence of such coarse-graining is the dramatic simplification of the energy landscape (Figure 1E). Due to the drastic reduction in the number of degrees of freedom (Figure 1D-E), and the relatively smooth topology of the coarse-grained energy landscape, studying processes over biologically relevant length and time-scales becomes far more tractable. Coarse-graining can exploit a ‘top-down’, ‘bottom-up’, or a combination of both approaches.^27^ Irrespective of the details, any systematic coarse-graining should only remove unnecessary ripples from the landscape and preserve the underlying physics (See Figure 1D and 1E). However, this is not always guaranteed.^28^ In recent years, various coarse-graining schemes for DNA have been developed. Schatz and coworkers introduced a two-bead DNA model to probe the denaturation dynamics of double-stranded DNA, but due to the exclusion of electrostatic interactions, salt dependence could not be explored.^29,30^ Subsequently, many higher resolution coarse-grained models have been proposed. The NARES models of Scheraga and coworkers^31–33^ have been used in a wide range of applications, from structure prediction^34^ to simulating DNA helix assembly.^32^ The HIRE-DNA model, which represents each nucleotide by either six or seven interaction sites, has been successfully employed to investigate the thermodynamics as well as the kinetics of DNA folding. ^35,36^ Owing to its unique ability to describe Hoogsteen interactions, HIRE-DNA has proven particularly well-suited to probe energy landscapes underlying G-quadruplex polymorphism.^37^ A number of studies have also employed the SIRAH force field,^38,39^ which has a resolution similar to HIRE-DNA, to study bubble formation^40^ and ion-induced bending of DNA.^41^ Surprisingly, the MARTINI framework, a trend-setter in CG simulations of proteins and lipids, has had limited success in the case of DNA. A recent study^42^ revealed that it cannot faithfully describe single-stranded DNA, or the stability of the DNA double helix unless an auxially elastic network is included to penalize deviations from the equilibrium structure.

Many CG models of DNA adopt the three interaction site (TIS) representation, originally formulated by Hyeon and Thirumalai^43^ The TIS resolution provides an ideal trade-off between accuracy and speed, and over the years has revealed many key aspects of DNA biophysics. In a series of papers, de Pablo and coworkers have used their 3SPN family of models^44–46^ to provide crucial insights into sequence-dependent DNA mechanics, bending fluctuations, as well as strand hybridization. The oxDNA framework,^47,48^ which also exploits the TIS mapping scheme, has been used to probe the free energy landscapes associated with diverse processes, including DNA self-assembly, hairpin folding, duplex bending, and conformational switching in DNA origami.^49^ Recently, Chakraborty *et al* introduced the Three Interaction Site (TIS) model for DNA (hereafter referred to as TIS-DNA) to provide a balanced description of both single-stranded and double-stranded DNA in terms of sequence-dependent thermodynamic and mechanical properties.^50^ Subsequently, TIS-DNA was integrated into COFFEE,^51^ a versatile framework for exploring the structure and dynamics of DNA-protein complexes, particularly nucleosomes. To probe genome organization at longer length scales, one needs to resort to more reductionist approaches, based on the 1CPN,^52^ or the CGeNArate models.^53^ In Table 1, we provide a representative overview of CG models that have been used to explore diverse problems in DNA biophysics. We emphasize that this list is merely illustrative and not meant to be exhaustive. For a more detailed discussion on various CG models of DNA, we refer the reader to a recent review by Pantano and coworkers.^54^

**Table 1:** Some commonly used coarse-grained (CG) models, and their applications in DNA biophysics.

|  | Model | Resolution per nucleotide | Applications |
| --- | --- | --- | --- |
| 1 | 1CPN <sup>52</sup> | 1 bead | Chromatin folding and organization. |
| 2 | Marenduzzo *et al.* <sup>25</sup> | 1 bead | DNA denaturation and supercoiling. |
| 3 | CGeNArate <sup>53</sup> | 1 bead | DNA supercoiling, Dynamics of yeast gene and the entire human mitochondrial DNA on the μs time-scale. |
| 4 | Schatz *et al.* <sup>29,30</sup> | 2 beads | DNA strand hybridization. |
| 5 | 3SPN <sup>44–46</sup> | 3 beads | DNA assembly, sequence-dependent thermodynamics and mechanics of dsDNA, nucleosome structure and dynamics. |
| 6 | oxDNA <sup>47–49</sup> | 3 beads | Self-assembly of DNA, force-extension curves, DNA nanotechnology, hairpin folding and origami landscapes. |
| 7 | TIS-DNA <sup>50,51</sup> | 3 beads | Hairpin folding, mechanical properties of ssDNA and dsDNA, nucleosome structure and dynamics. |
| 8 | NARES <sup>31–34</sup> | 3 beads | Structure-prediction, DNA helix assembly, dynamics of protein-DNA complexes. |
| 9 | HIRE-DNA <sup>35–37</sup> | 6 to 7 beads | Thermodynamics and kinetics of DNA folding, Energy landscape analysis of G-quadruplex polymorphism. |
| 10 | SIRAH <sup>38–41</sup> | 6 beads | DNA bubble formation, Ion-induced deformation of DNA double helix, tetranucleosome dynamics. |
| 11 | MARTINI <sup>42</sup> | 6 to 7 beads | Mapping counter-ion distributions around dsDNA, dynamics of DNA oligomers. |

In this chapter, we illustrate how the TIS-DNA model can be employed to explore the folding landscape of a prototypical DNA tetraloop hairpin. The remainder of the chapter is organized as follows: First, we provide a brief description of the TIS-DNA model, specifying only the key aspects of its parametrization process. Detailed information, including the force-field parameters, can be found in the original report.^50^ Next, we present our results on the thermodynamics and kinetics of hairpin folding, along with a discussion of the various algorithms used for landscape exploration. Finally, we outline some current challenges, and highlight possible avenues for future exploration.

## Energy landscape exploration using the Three Interaction Site (TIS) model of DNA

### The TIS-DNA Model

The Three Interaction Site (TIS) model of DNA^50^ (referred to as TIS-DNA) represents each nucleotide by three interaction sites. In this representation, spherical beads are placed at the centers of masses of the phosphate group, the sugar ring and the nucleobase (see Figure 1A). The potential energy function has the same form as the TIS-RNA model of Denesyuk and Thirumalai. ^55^ The potential energy, *U*_*T IS*−*DNA*_, includes contributions from bonded (*U*_*B*_), angular (*U*_*A*_), single-stranded stacking (*U*_*S*_), hydrogen-bonding (*U*_*HB*_), excluded volume (*U*_*EV*_) and electrostatic (*U*_*ELE*_) interactions:

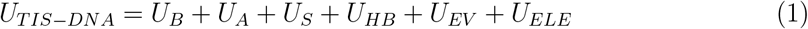

Harmonic potentials are used to describe the bonded and angular interactions:

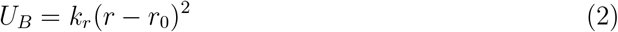

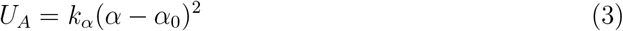

In Equations 2 and 3, *r*_0_ and *α*_0_ denote the equilibrium values of the bond length and bond angle, respectively, and *k*_*r*_ and *k*_*α*_ denote the corresponding force constants. The optimal values of *k*_*r*_ and *k*_*α*_ were determined using a Boltzmann Inversion procedure.^50^

To prevent unphysical overlap among beads, we use the Weeks-Chandler-Andersen (WCA) potential^56^ to denote the non-covalent interactions:

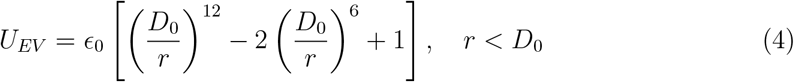

Although the choice of the WCA potential is by no means unique, ^57^ it ensures computational efficiency, as the excluded volume term vanishes when the distance beween the two beads is greater than *D*_0_.

#### Stacking and hydrogen-bonding interactions

Stacking interactions along with hydrogen-bonding between complementary strands are crucial for the stability of the DNA duplex. By directly measuring stacking free energies, Frank-Kamenetskii and coworkers established that DNA stability at different temperatures and salt concentrations is largely dictated by base-stacking. ^58^ In the TIS-DNA model, only stacking interactions between consecutive nucleotides along a DNA strand is considered, and is given by:

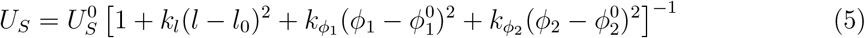

In Equation 5, 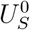, sets the energy scale for base-stacking, and for each dimer it was calibrated to reproduce the nearest-nearest-neighbor thermodynamics, as described by Santalucia and Hicks.^59^ Deviations from the equilibrium geometry (described by stacking distance, *l*_0_, and backbone dihedrals, 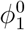 and 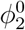) influence the strength of stacking interactions. The equilibrium stacking distance of each dimer and the corresponding backbone dihedrals were determined by coarse-graining an idealized B-DNA helix to the TIS resolution. The force constants, *k*_*l*_, 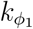 and 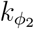 were determined from a Boltzmann inversion of the distributions of the stacking distances and dihedrals computed from a database of experimental structures.^50^ In the original implementation of the TIS-DNA model, only canonical hydrogen-bonding interactions (Watson-Crick) were included, although extensions to incorporate other base pairing modes are certainly possible.^60^ The potential energy function describing hydrogen-bonding interactions is given by:

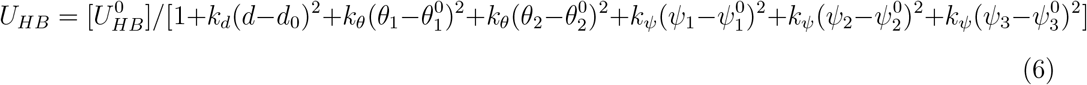

In Equation 6, *U*_*HB*_ describes a single hydrogen bond. It is multiplied by either 2 or 3, depending on the interacting nucleotides (A-T or G-C). The strength of hydrogen-bonding, denoted by 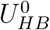, was optimized to reproduce the experimental melting profile of a DNA hairpin.^50^ Similar to base-stacking interactions, hydrogen-bonding potential is modulated by deviations from an idealized B-DNA helical geometry, quantified in terms of the internucleotide distance *d*, angles *θ*_1_ and *θ*_2_ and dihedrals *ψ*_1_, *ψ*_2_ and *ψ*_3_. The parameters *k*_*d*_, *k*_*θ*_, and *k*_*ψ*_ were also determined from a Boltzmann inversion procedure like the other harmonic force constants. In Figure 2, we provide a schematic description of the geometric parameters in terms of which the stacking and hydrogen-bonding interactions are represented within the TIS-DNA model.

**Figure 2.**
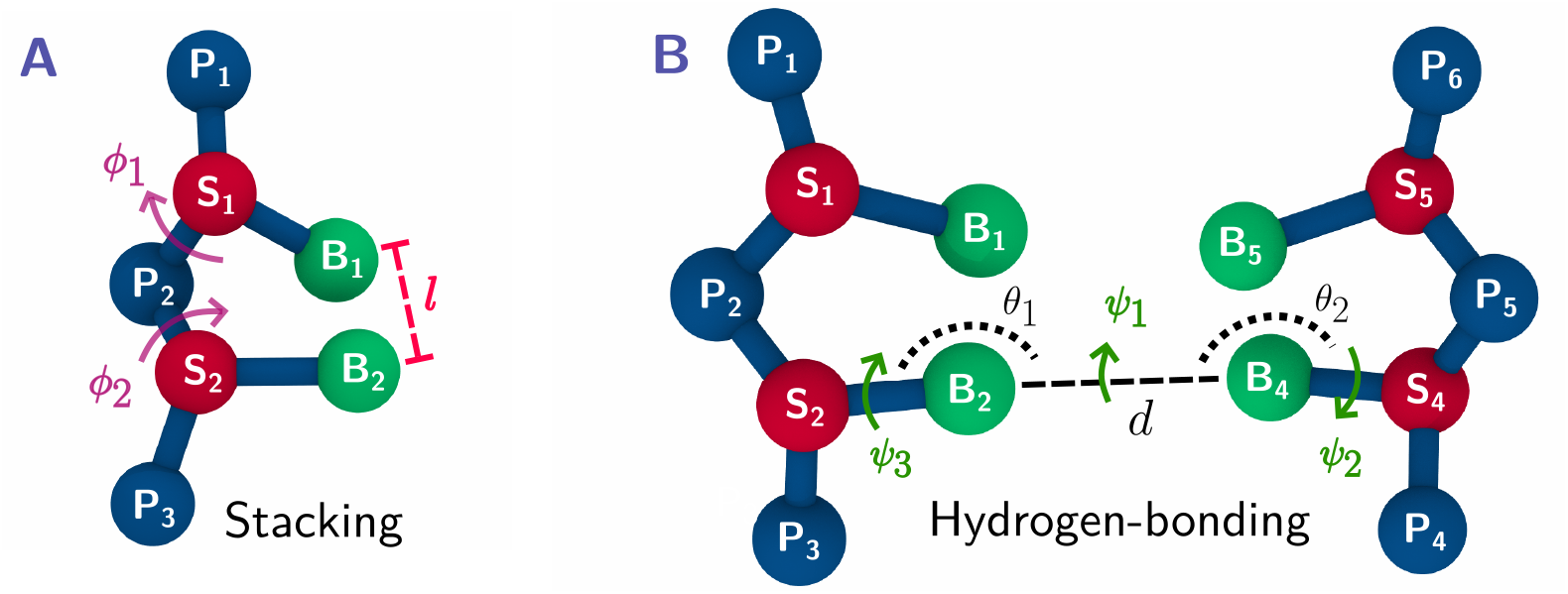
Illustration of (A) the distance *l* and backbone dihedrals, *φ*_1_ and *φ*_2_ used for defining the base-stacking interaction in equation 5). (B) distance, *d*, angles *θ*_1_ and *θ*_2_ and dihderals *ψ*_1_, *ψ*_2_ and *ψ*_3_, which control the strength of the hydrogen-bonding interaction (equation 6)

#### Electrostatic interactions

The electrostatic interactions between phosphate beads are described implicitly using the Debye-Hückel theory: ^61^

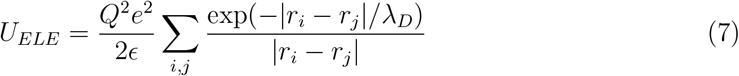

where |*r*_*i*_ *− r*_*j*_| denotes the separation between two phosphate beads *i* and *j*; *ϵ* is the dielectric constant of water; and *λ*_*D*_ is the Debye length, which describes how monovalent salt attenuates electrostatic interactions.

In the spirit of Oosawa-Manning counterion condensation,^62^ we use a renormalized charge on the phosphate beads given by:

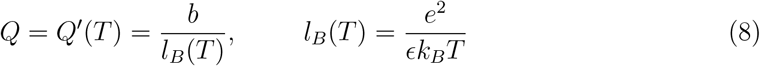

In Equation 8, *b* denotes the length per unit charge (set to 4.0 Å following Olson and coworkers^63^); and *l*_*B*_ is the Bjerrum length, the distance at which the electrostatic energy becomes comparable to the thermal energy, *k*_*B*_*T*. The temperature dependence of *Q* is nonlinear as the dielectric constant *E* itself varies with temperature. We use the well-known equation of Malmberg and Maryott^64^ that accurately fits experimental measurements of the dielectric constants of water over a wide temperature range.

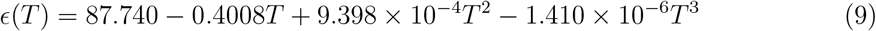

In Equation 9, *T* denotes the temperature in Celsius. While an implicit description of electrostatics may be reasonable for many applications in DNA biophysics, it is insufficient to describe the effect of divalent (or multivalent) ions on the structure and folding dynamics. Recently, an extension to TIS-DNA was proposed,^65^ in which various counterions, such as Na^+^, Mg^2+^ and Ca^2+^ were included explicitly. Interestingly, this new model (christened TIS-ION) works entirely “out of the box” without requiring any further fine-tuning of the TIS-DNA force field parameters.

### DNA Hairpin folding landscape

In principle, the TIS-DNA model is well-suited for probing rather complex energy landscapes, including those of DNA pseudoknots, Holliday junctions or cruciforms. However, to streamline the discussion in the remainder of this chapter, we demonstrate its applicability to a tetraloop DNA hairpin, having the sequence 5-^t^GGATAA(T_4_)TTATCC-3^t^. We directly compare our results to the work of Ansari and coworkers, who studied the folding of this DNA hairpin using both fluorescence spectroscopy and laser-induced temperature jump experiments.^66^ A schematic of the hairpin structure, along with its sequence is shown in Figure 3A.

**Figure 3.**
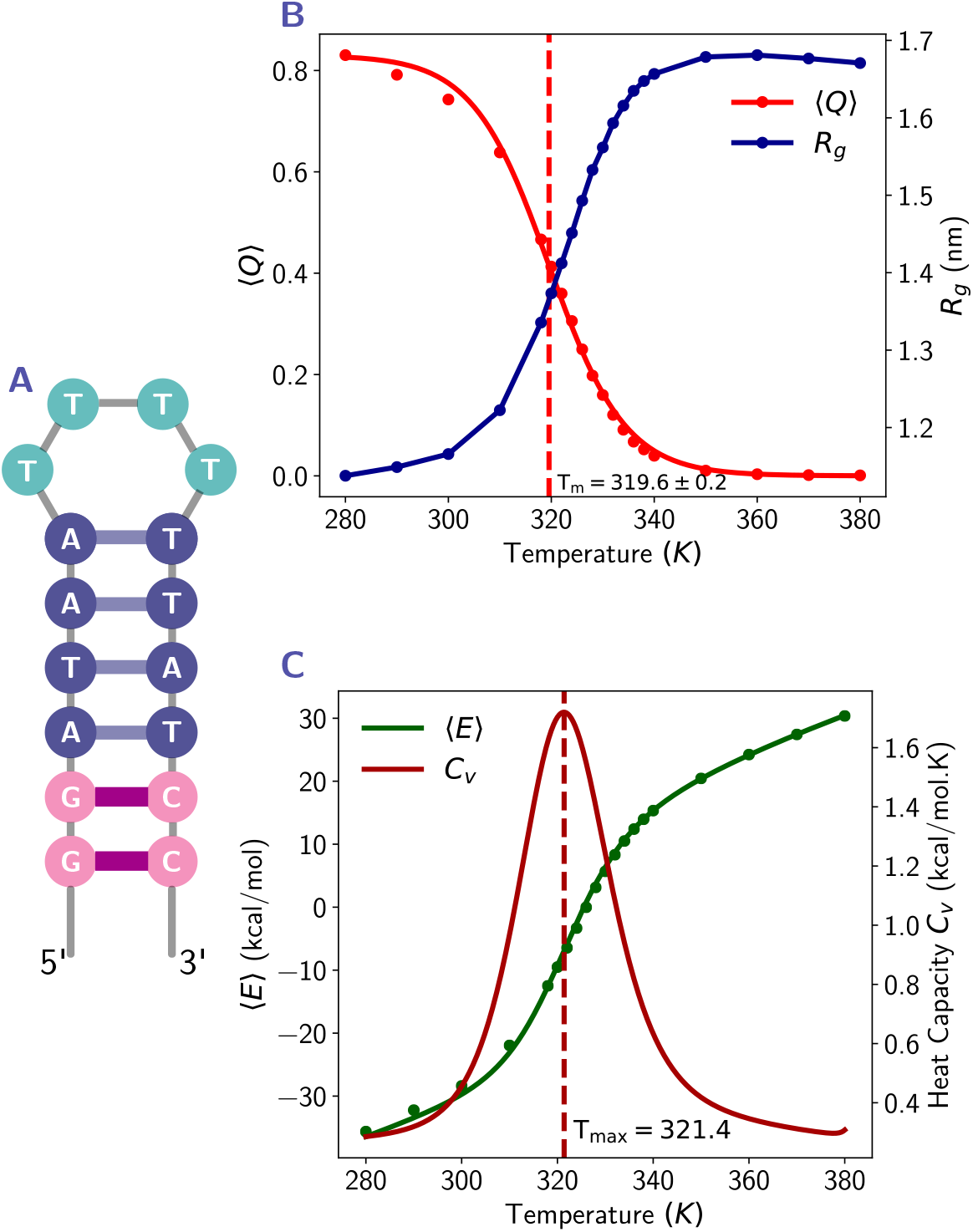
(A) Schematic of the DNA Hairpin considered in this study. This folding of this hairpin sequence has been investigated by Ansari and coworkers^66^ using a combination of laser-induced temperature jump and flourescence spectroscopy experiments. (B) The variation of *R*_*g*_ (blue circles) and fraction of native contacts (red circles), ⟨*Q*⟩ with temperature. By fitting the temperature-dependence of ⟨*Q*⟩ to a sigmodial logistic function (shown as a solid red curve), we determine the melting temperature, *T*_*m*_ to be ≈ 319 K. (C) The variation of the potential energy, ⟨*E*⟩ (green circles) and heat capacity, *C*_*v*_ (maroon curve) with temperature. The heat capacity profile exhibits a peak at *T*_*max*_ = 321 K, which is close to the *T*_*m*_ determined from equation 12.

#### Folding thermodynamics

To explore the folding thermodynamics, we carried out Lanvgevin dynamics simulations in the low-friction regime. The trajectories were generated by integrating the following equation of motion for each bead

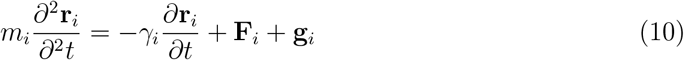

In Equation 10, **F**_*i*_ denotes the conservative force derived from the TIS-DNA potential energy function, while **g**_*i*_ is a Gaussian random force acting on bead *i* having mass *m*_*i*_. The viscous drag on each bead is given by *γ*_*i*_. It is computed using Stokes’ law, *γ*_*i*_ = 6*πηR*_*i*_, where *η* is the viscosity of the surrounding medium and *R*_*i*_ is the hydrodynamic radius. To enhance conformational sampling, we chose a value of 10^−5^ Pa.s for *η*, which is about 1% of the viscosity of water. It is important to note that the particular choice of *η* has no bearing on thermodynamic properties. The random force, **g**_*i*_ and the *γ*_*i*_ are related through the fluctuation dissipation theorem, ⟨**g**_*i*_(*t*)**g**_*j*_(*t*^t^)⟩ = 6*k*_*B*_*Tγ*_*i*_*δ*_*ij*_*δ*(*t−t*^t^), ensuring that the sampling follows the correct canonical distribution. The values for *R*_*i*_ are assigned as follows: 2.0 Å for the phosphate beads, 2.9 Å for the sugar beads, 3.0 Å for the guanine beads, 2.8 Å for adenine beads, and 2.7 Å for the cytosine and thymine beads. In the TIS-DNA model, the average mass *m* of a bead is about 116 g/mol, the typical length scale, *a* is ≈ 3.9 Å and the energy scale, *ϵ* is 1.0 kcal/mol. From these values, we estimate the natural unit of time in the low-friction regime to be 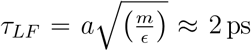. We used a time-step of 0.10 τ_*LF*_ to integrate the equation of motion (Equation 10) employing a modified version of the velocity-Verlet scheme described by Honeycutt and Thirumalai.^67^ To generate sufficient statistics for the thermodynamic observables, simulations were carried out for 2 *×* 10^9^ steps at each temperature starting from ten different initial conditions.

We characterize the folding thermodynamics using two structural order parameters, the fraction of native contacts, ⟨*Q*⟩, and the radius of gyration, *R*_*g*_. These are described below:

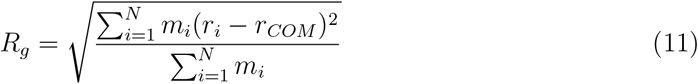

In Equation 11, *r*_*i*_ and *m*_*i*_ denote the position and mass of the *i*^*th*^ bead, respectively, and *r*_*COM*_ denotes the center of mass. The radius of gyration, *R*_*g*_ describes the global dimensions of the DNA chain, and can be experimentally determined from small-angle scattering(x-ray and neutron). In recent times, small-angle x-ray scattering(SAXS) has emerged as a potent tool for interrogating the folding landscape of nucleic acids.^68^

In addition to *R*_*g*_, the fraction of native contacts ⟨*Q*⟩ is a commonly used order parameter in folding studies.^22,69^ Experimentally, the measurable quantity is typically 1 *−* ⟨*Q*⟩, which is related to the changes in relative absorbance. From our simulations, we compute ⟨*Q*⟩ by assuming that a hydrogen-bond is formed if *U*_*HB*_ *< nk*_*B*_*T*, with *n* = 2 for a A-T base pair and *n* = 3 for a G-C base pair.

In Figure 3B we show the variation of ⟨*Q*⟩ and *R*_*g*_ with temperature. As expected, the hairpin unfolds with increasing temperature. There is a gradual reduction in the number of native contacts, and a concomitant increase in chain dimensions. We estimate the melting temperature, *T*_*m*_ by fitting the variation of ⟨*Q*⟩ to the following sigmoidal curve:

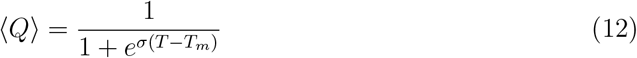

From the fit we obtain *T*_*m*_ ≈ 320 K, which is in quantitative agreement with the value reported from by Ansari and coworkers.^66^ In equation 12, *σ* denotes the width (or the cooperativity) of the melting transition, and as compared to experiment, it is usually over-estimated. This disparity likely arises from the omission of non-native interactions in the TIS-DNA force field, as well as the use of isotropic potentials for describing the excluded volume interactions.

Another key thermodynamic variable, the heat capacity, *C*_*v*_, was determined from the fluctuations of potential energy, *E*, at each temperature:

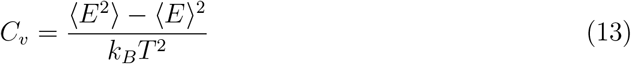

We combined data from different temperatures using the weighted histogram analysis method (WHAM)^70^ to generate a smooth heat capacity profile. As illustrated in Figure 3C, the average potential energy, ⟨*E*⟩ increases with temperature, implying that high-lying local minima on the energy landscape become progressively accessible and contribute to the global thermodynamics. The potential energy fluctuations are enhanced in the vicinity of the melting temperature, and a pronounced peak appears in the heat capacity curve at *T*_*max*_ ≈ 321 K (Figure 3C), which is close to the *T*_*m*_ we determine from equation 12. The various thermodynamic observables exhibit the hallmarks of a phase transition in a finite system,^71^ consistent with a two-state model.

#### Kinetics of folding

We carried out Brownian dynamics simulations in the high friction regime (corresponding to a solvent viscosity of 10^−3^ Pa.s) to probe the folding kinetics. The motion of each bead *i* is given by:

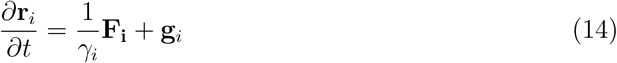

In this regime, the natural unit of time is given by 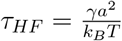. Using the values appropriate for the TIS-DNA model, we obtain τ_*HF*_ ≈ 262 ps. We employed a time-step of 0.1τ_*HF*_ to integrate the equation of motion using the Ermak-McCammon algorithm.^72^ Simulations were initiated from one hundred distinct initial conditions at 300 K to obtain a converged free energy surface (FES), and a reliable estimate of the folding time.

A two-dimensional representation of the free energy landscape in terms of pre-defined order parameters, *q*_1_ and *q*_2_ can be obtained from their joint probability distribution *P* (*q*_1_, *q*_2_) as follows:

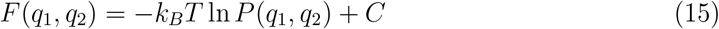

where *C* is an arbitrary constant, and *q*_1_ and *q*_2_ can represent structural metrics, such as *R*_*g*_, root-mean-squared-deviation (RMSD), number of native contacts, or other complex collective variables that are commonly used as proxies for the “reaction coordinate”.

The folding landscape of the DNA hairpin in terms of the radius of gyration, *R*_*g*_ and a dimensionless quantity, 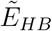 is shown in Figure 4A. Specifically, 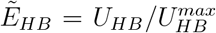, defined in terms of the hydrogen-bonding energy, serves as measure of the native contact formation, while *R*_*g*_ describes the global dimension of the DNA chain. Although 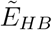 is highly correlated to ⟨*q*⟩, we choose the former as it brings out the underlying features of the free energy landscape, particularly the ‘funnel-like’ character, more effectively. The native basin (centered at *R*_*g*_ ≈ 1.1 nm and 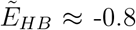) consists of folded hairpins. In these structures (snapshot 2), all nucleobases except the terminal ones engage in Watson-Crick interactions. This behavior of the terminal base-pair, often referred to as “fraying-peeling”, has also been reported previously in atomistic simulations.^73^ Near-native structures, which exhibit a well-formed loop region, but lack some of the base-pairing interactions (snapshot 3), are destabilized with respect to the folded state by about 2.7 kcal/mol. Relatively compact unfolded states (snapshot 1) lie about 1.5 kcal/mol above the folded state. These structures exhibit different extents of ‘non-specific’ collapse, with some configurations having opposite strands aligned in a manner that promotes folding. Fully extended states are destabilized by about 4.7, kcal/mol relative to the native state, and have low equilibrium population at 300 K. The organization of the various structural ensembles on the free energy landscape provides important insight into the plausible folding mechanism. In the first stage, the chain is likely to undergo non-specific collapse that bring the two opposing strands in contact. Subsequently, the loop becomes more structured and various contacts are established in the hairpin stem. The final stage of folding involves rapid zippering of the hairpin stem to form the native state. Due to the neglect of non-native interactions in the TIS-DNA model, the mechanism may appear somewhat simplistic; nevertheless, it recapitulates many features of previously proposed folding pathways.^22,74,75^

**Figure 4.**
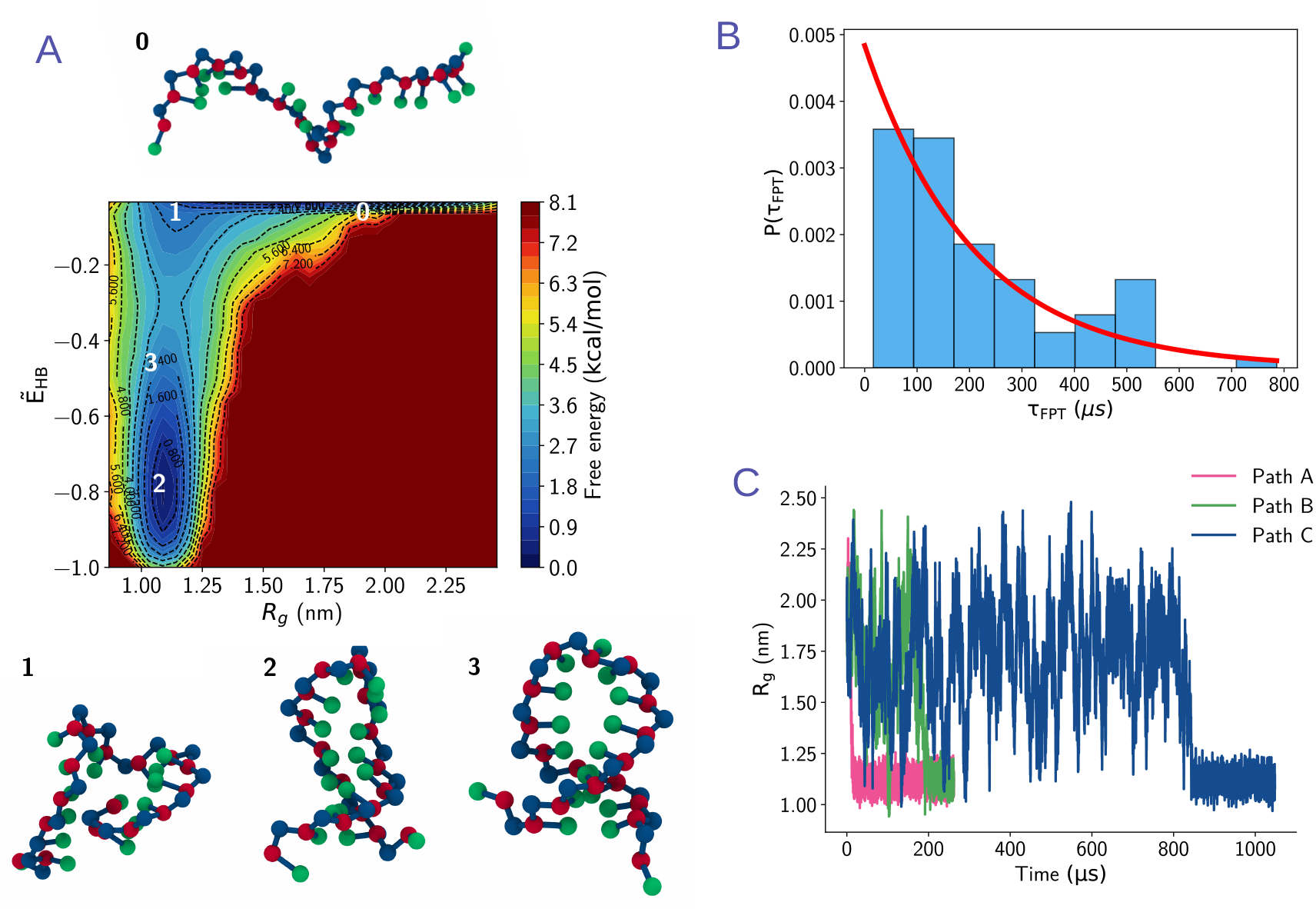
(A) A two-dimensional free energy landscape for the DNA hairpin. The two order parameters, *R*_*g*_ and 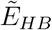 give complementary insights into the folding process. While *R*_*g*_ characterizes the global chain dimension, 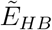 quantifies the number of native contacts present in a given configuration. A few representative snapshots are also shown superimposed on the landscape: folded hairpin (snapshot 2), near-native state with some base-pairs missing (snapshot 3), compact structure without any native contacts (snapshot 1) and the fully extended state (snapshot 0). (B) Distribution of the first passage times for reaching the native hairpin configuration starting from a fully extended state. The solid red line is a fit to the Poisson distribution. The MFPT, ⟨τ_*F PT*_⟩, is estimated to be ≈206 *µ*s.(C) Some typical folding trajectories with *R*_*g*_ as a progress variable. Path A corresponds to a fast pathway associated with a folding time of τ_*F*_≈ 16 *µ*s, path B is associated with a τ_*F*_ of 200 *µ*s ≈ ⟨*τ*_*F PT*_⟩, path C is the slowest and is characterized by the DNA chain undergoing multiple excursions on the energy landscape until the two opposing strands are approximately aligned to nucleate the first native contact.

To quantify the folding kinetics, we calculated the distribution of the first passage times (FPTs) for reaching the native state. Mathematically, it is defined as:

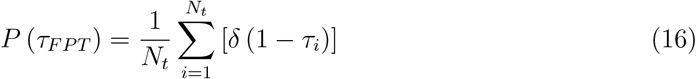

where *N*_*t*_ denotes the number of independent trajectories, and τ_*i*_ denotes the time in the *i*^*th*^ trajectory when the folded basin is visited for the first time. Guided by the topography of the free energy landscape, we define a folded state as any structure having *R*_*g*_ ≤ 1.2 Å, and 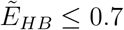.

The mean first passage time (MFPT) for the folding process, τ_*F PT*_, initiated from the fully extended state is estimated to be ≈ 206 *µ*s. Our results are consistent with the previous simulations of Hyeon and Thirumalai,^43^ as well as the experimental work of Ansari and coworkers.^4^ The FPT distribution (Figure 4B) is approximately Poissonian in shape, indicating that transitions to the hairpin structure do not involve any long-lived metastable states. The width of the distribution further suggests that folding occurs via multiple routes. In Figure 4C, we depict some representative folding pathways, using *R*_*g*_ as a progress variable. The hairpin folds the fastest along path A (folding time, τ_*F*_ ≈ 16 *µ*s), in which the initial collapse itself aligns the opposing strands in a near-perfect orientation, allowing native contacts to form rapidly in a ‘downhill’ fashion. Along path B (τ_*F*_ *≈ τ*_*F PT*_ ≈ 200 *µ*s), there are multiple stages of chain collapse and expansion before base-pairing interactions are formed. Folding is the slowest along path C (τ_*F*_ ≈ 800 *µ*s) and the *R*_*g*_ fluctuates substantially, prior to the emergence of any native-like contacts. A closer inspection of the folding trajectories reveals that G2–C17 is the first base-pairing interaction to form along path A, while A4–T15 is the earliest to appear along path C. Overall, the folding mechanisms seem consistent with the “compaction” model proposed by Pande and coworkers.^74^

### Current challenges and future outlook

Compared to models of similar resolution, TIS-DNA provides a balanced description of the sequence-dependent properties of both single-stranded and double-stranded DNA.^50^ In this chapter, we have demonstrated how it provides a quantitative description of hairpin folding thermodynamics and captures several key features of the folding kinetics. Despite its simplicity, we envision that TIS-DNA would be particularly suitable for providing critical insights into diverse problems at the interfaces of physics and biology. For instance, the salt and force-induced reshaping of DNA energy landscapes is not fully resolved, and we believe that the TIS-DNA model is well-poised to address some of the existing paradoxes.

The current version of TIS-DNA has several limitations. It cannot be used for structure-prediction *per se* (like NARES and other CG models) as it requires some prior information of the base-stacking and base-pairing interactions within the native state. This caveat, however, can be readily addressed as enumerating all possible base-stacking and base-pairing interactions for a given sequence amounts to little more than a book-keeping exercise. Indeed, work along these lines has already been reported by Denesyuk and Thirumalai in the context of the TIS-RNA model.^76^

Our study showcases TIS-DNA in its original incarnation, which does not have an explicit representation of electrostatics or solvation. As shown in previous studies,^74,77^ excluding such effects could give a misleading picture of the energy landscape, especially when folding is driven by ion-coordination or solvent-mediated interactions. The TIS-ION framework,^65^ addresses one of these limitations, by including the counter-ion atmosphere around DNA explicitly. Integrating water models in a seamless manner with existing CG representations of DNA, however, is not straightforward. Taking cues from the work of Molinero and coworkers,^78^ it would be possible to integrate various CG water models,^78^ such as monowater (mW) and big multipole water (BMW)^79^ into the TIS-DNA framework using a bottom-up approach, but one must remain cognizant of the potential mismatch in length scales.

The unprecedented breakthroughs in artificial intelligence (AI) and machine learning (ML) are poised to shape the future of force-field development, particularly coarse-graining endeavors. Various flavors of AI/ML have already made in-roads into existing pipelines,^80^ replacing or in some cases augmenting traditional physics-based approaches. We anticipate that future extensions of TIS-DNA will greatly benefit from these advances, and will be tailor-made for various applications in DNA biophysics.

## Acknowledgement

We thank the high performance computing (HPC) facility at The Institute of Mathematical Sciences for computing time. DC acknowledges the Anusandhan National Research Foundation (ANRF) for the Prime Minister’s Early Career Grant (ANRF/ECRG/2024/003762/CS). DC also acknowledges support via the sub-projec titled “Modeling of Soft Materials” within the IMSc Apex Project, funded by the Department of Atomic Energy, Government of India. KB thanks ANRF for a National Postdoctoral Fellowship (NPDF).

## Notes

### Competing Interest Statement

The authors have declared no competing interest.

